## Supplemental Figures for "Dengue after Zika: characterizing impacts of Zika emergence on endemic dengue transmission"

Borchering et al.

**SI Figure 1:** Dengue seasonality by biweeks with the top three highest incidence values for each season. Seasons are defined to range from biweek 18 to biweek 17 of the following year (biweek 26 is represented as biweek 0 below). Frequencies of peak biweek are displayed for each state in Brazil (**a**) and department in Colombia (**b**). Black lines show the average frequency for each biweek across locations within each region.

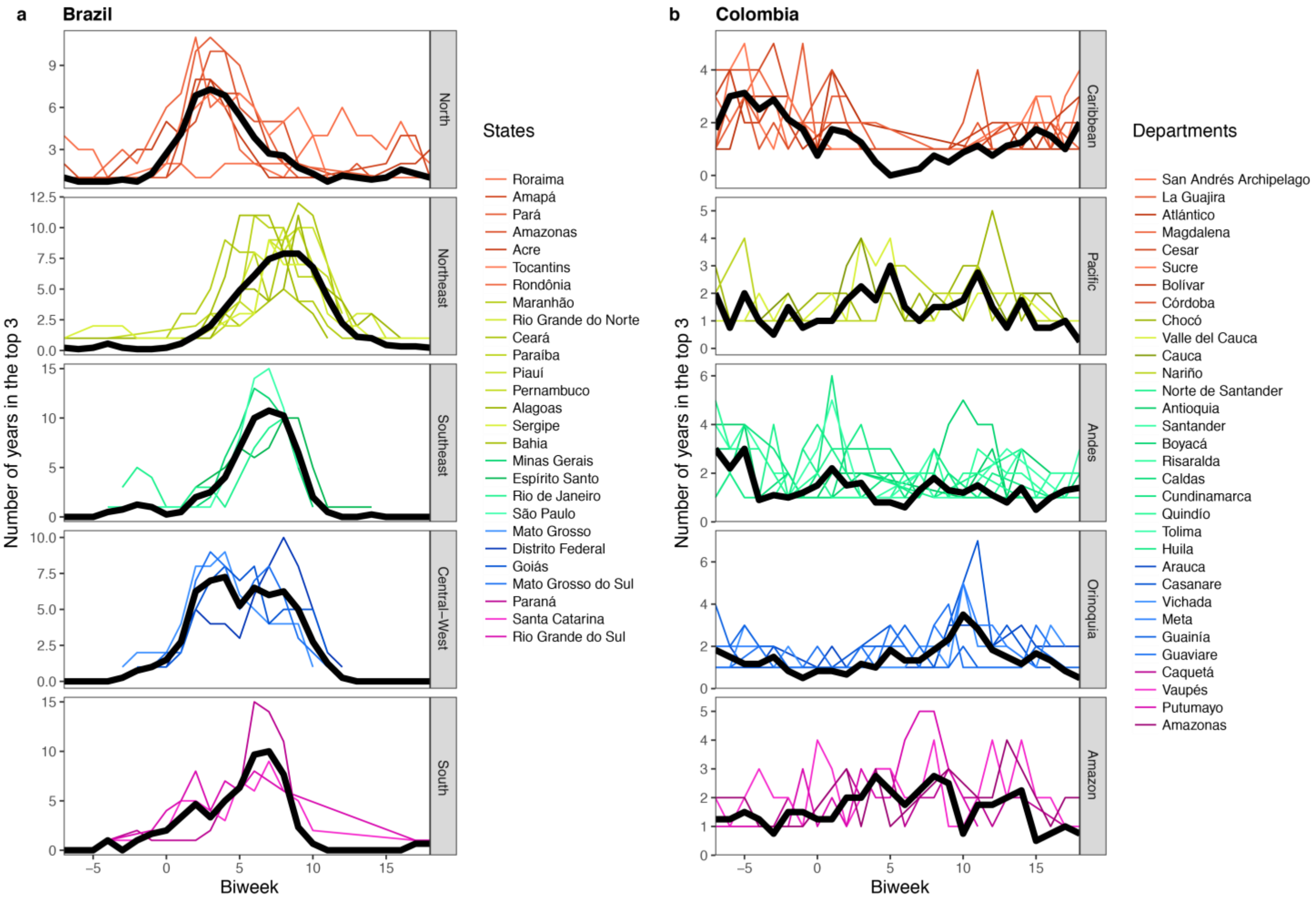

**SI Figure 2:** Variation in the stan model coefficients for each biweek. Results are shown for coefficients aggregated at the region (**a, c**) and state (Brazil in **b**) or department (Colombia in **d**) level.

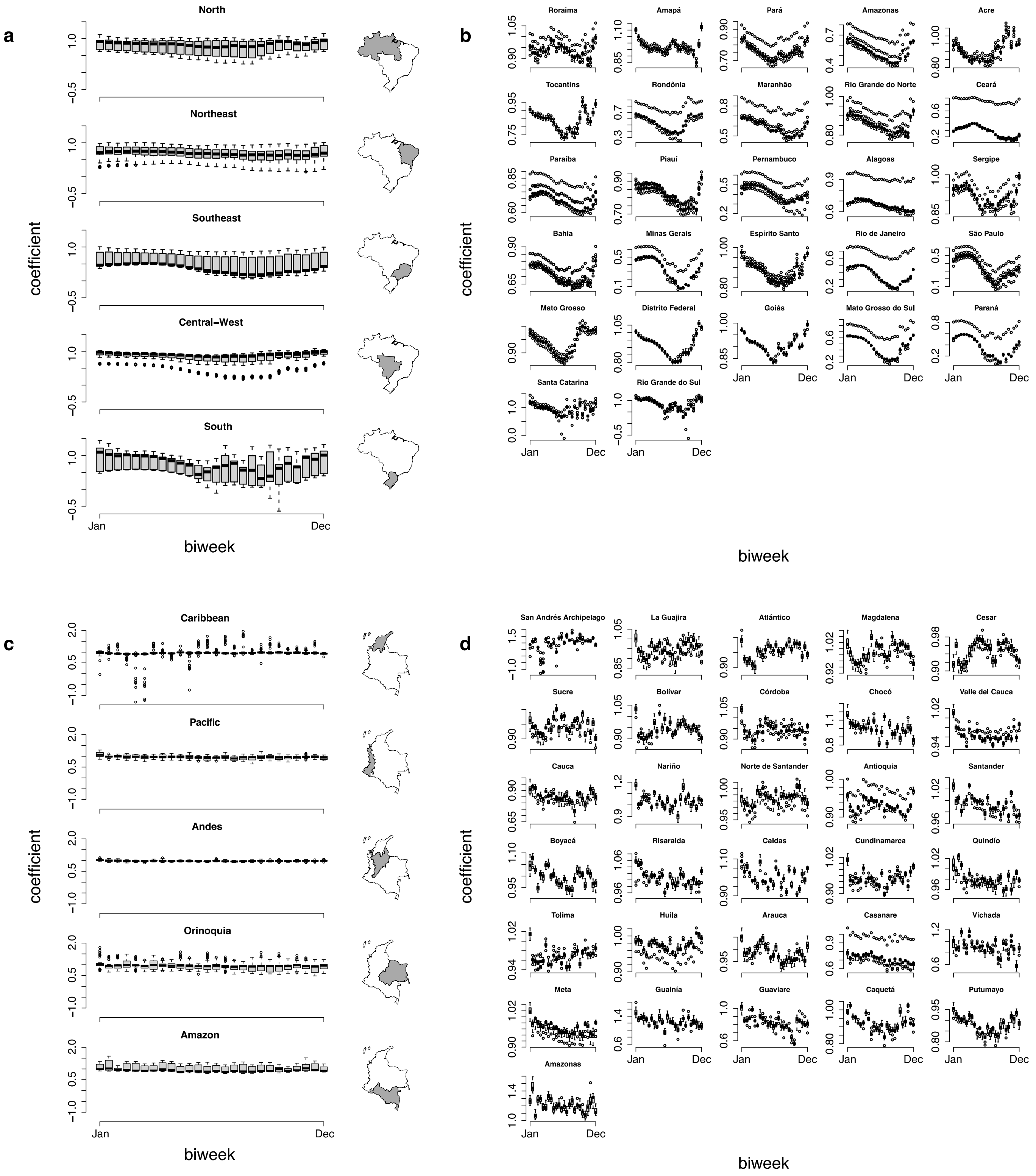

**SI Figure 3:** Bayesian R-squared plots for subnational location specific stan model predictions versus observed incidence for Brazil (**a**) and Colombia (**b**). The gray line corresponds the R-squared value for all of the biweeks that were included in the model fitting. The black dots show the Bayesian R-squared value for the year when it was left out of the model fitting. The x-axis indicates the year that was left out, and ranges over all years in the dataset 1999-2017 in Brazil and 2000-2017 in Colombia.

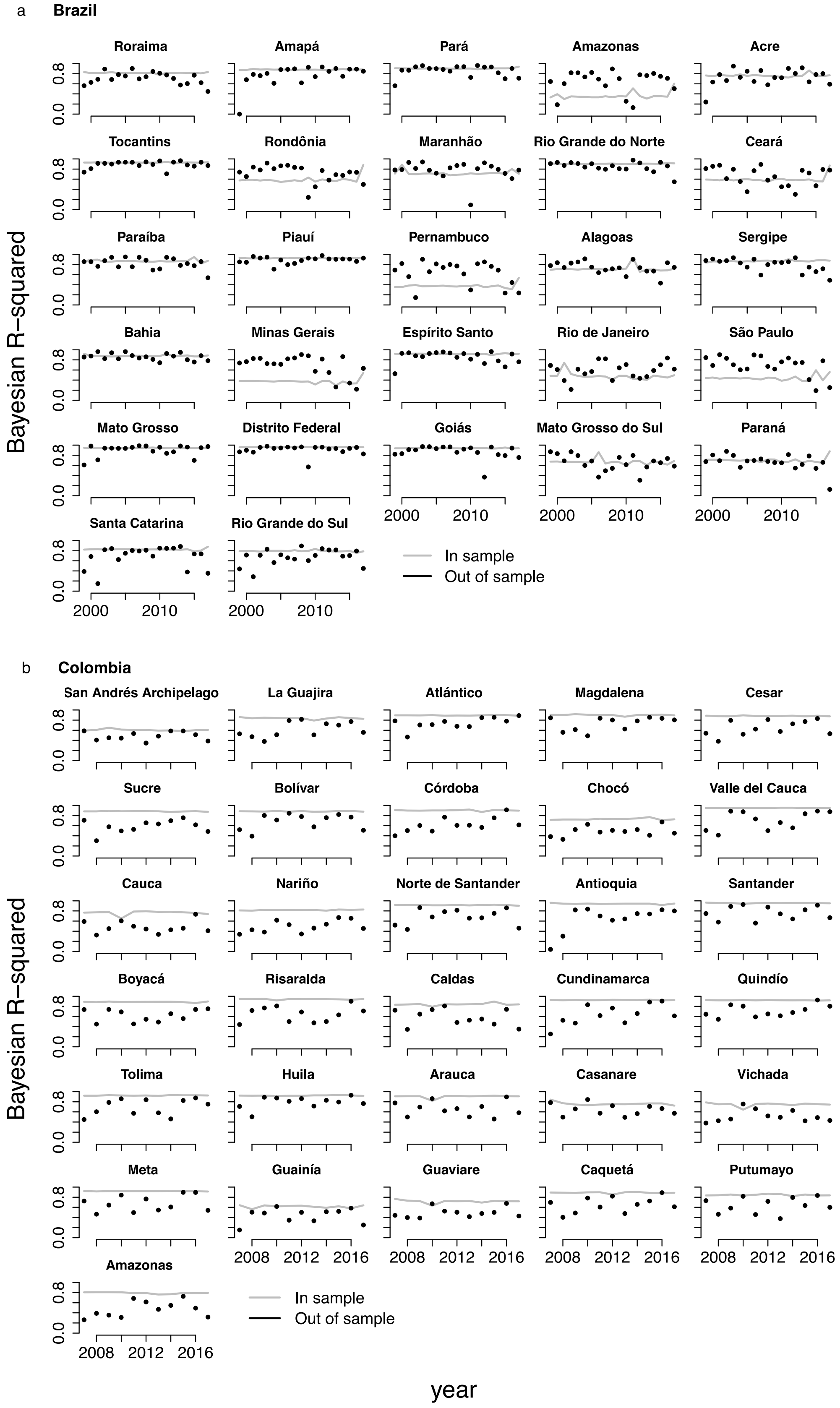

**SI Figure 4:** Comparison between predicted and observed incidence. Red and blue indicate that the observed incidence was above and below the median value of the posterior distribution of predicted values for that biweek, respectively. Dark biweeks indicate that the observed incidence was outside of the 95% prediction interval and medium shaded biweeks indicate that the observed incidence was outside of the 90% prediction interval. Results are shown for Brazil (**a**) and Colombia (**b**).

**a Brazil**

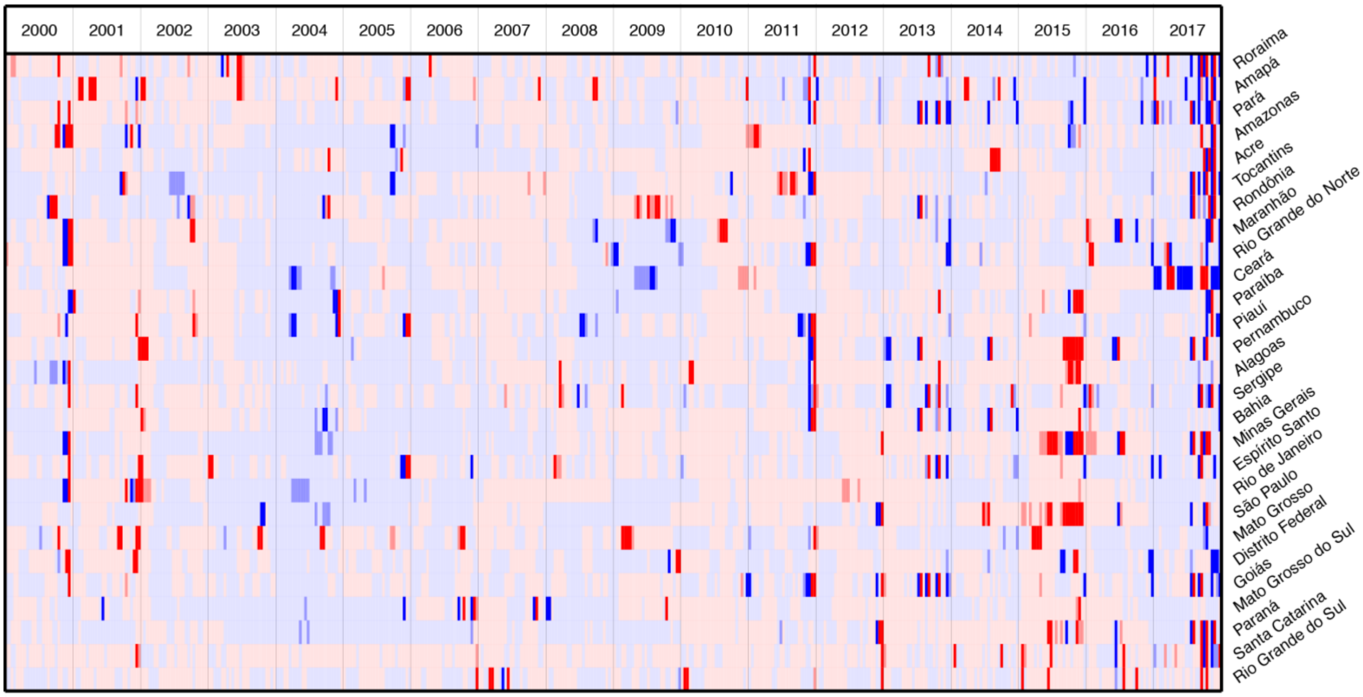


**b Colombia**

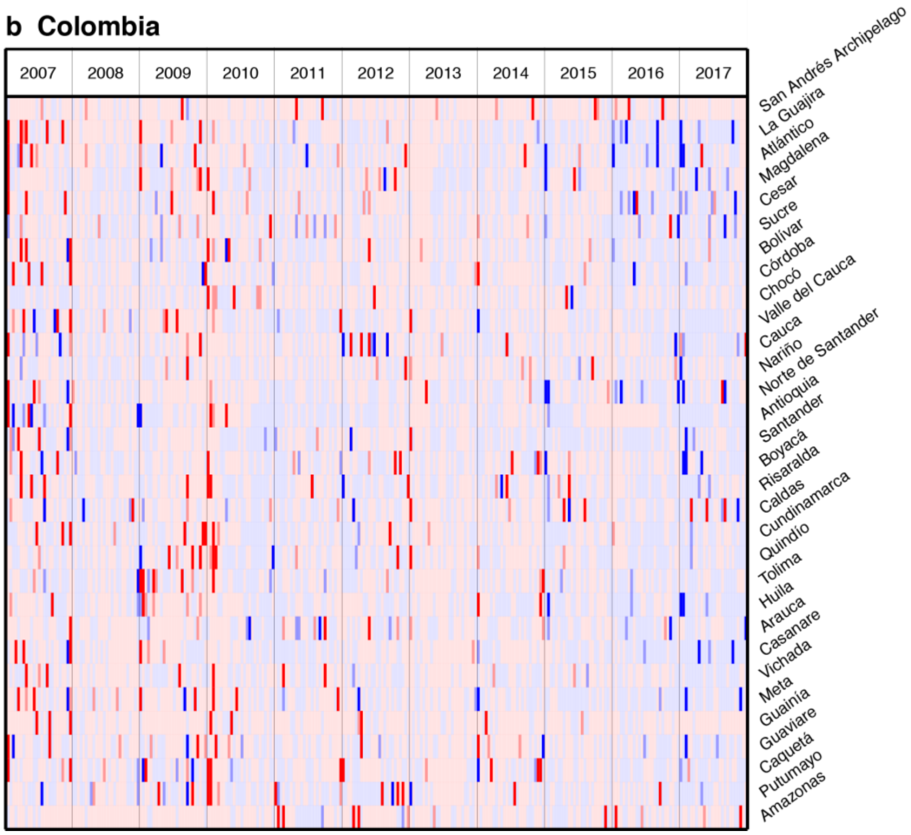

**SI Figure 5:** Bonferroni adjusted quantile plot for full time series and recent years. Quantile values are based on the location of the observed incidence in the cumulative distribution of 500 sampled posterior prediction values. Results are shown for Brazil (**a**) and Colombia (**b**).

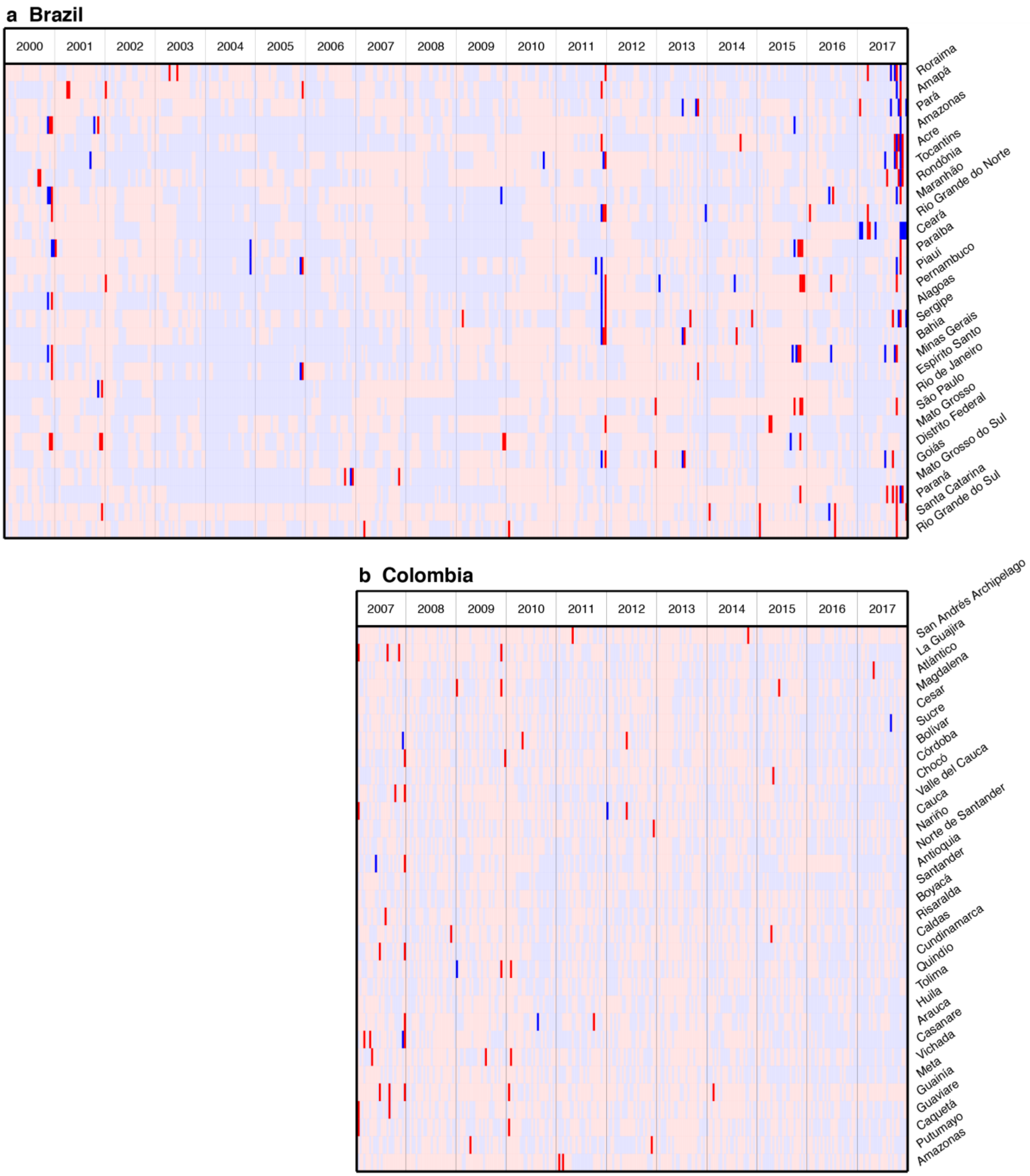

**SI Figure 6:** Dengue time series model results with spatial hierarchical structure and arboviral disease covariates. Mean and 95% Bayesian credible intervals are displayed for the shared effect (top row) and for the location specific effects (other rows ordered by region and then latitude) for Brazil (**a-d**) and Colombia (**e-h**). Location specific effects are displayed as the sum of shared coefficient and location specific coefficient. Models are displayed for biweekly Zika cases (**b**, **f**), biweekly chikungunya cases (**d**, **h**), cumulative Zika cases (**a, e**), and cumulative chikungunya cases (**c, g**).

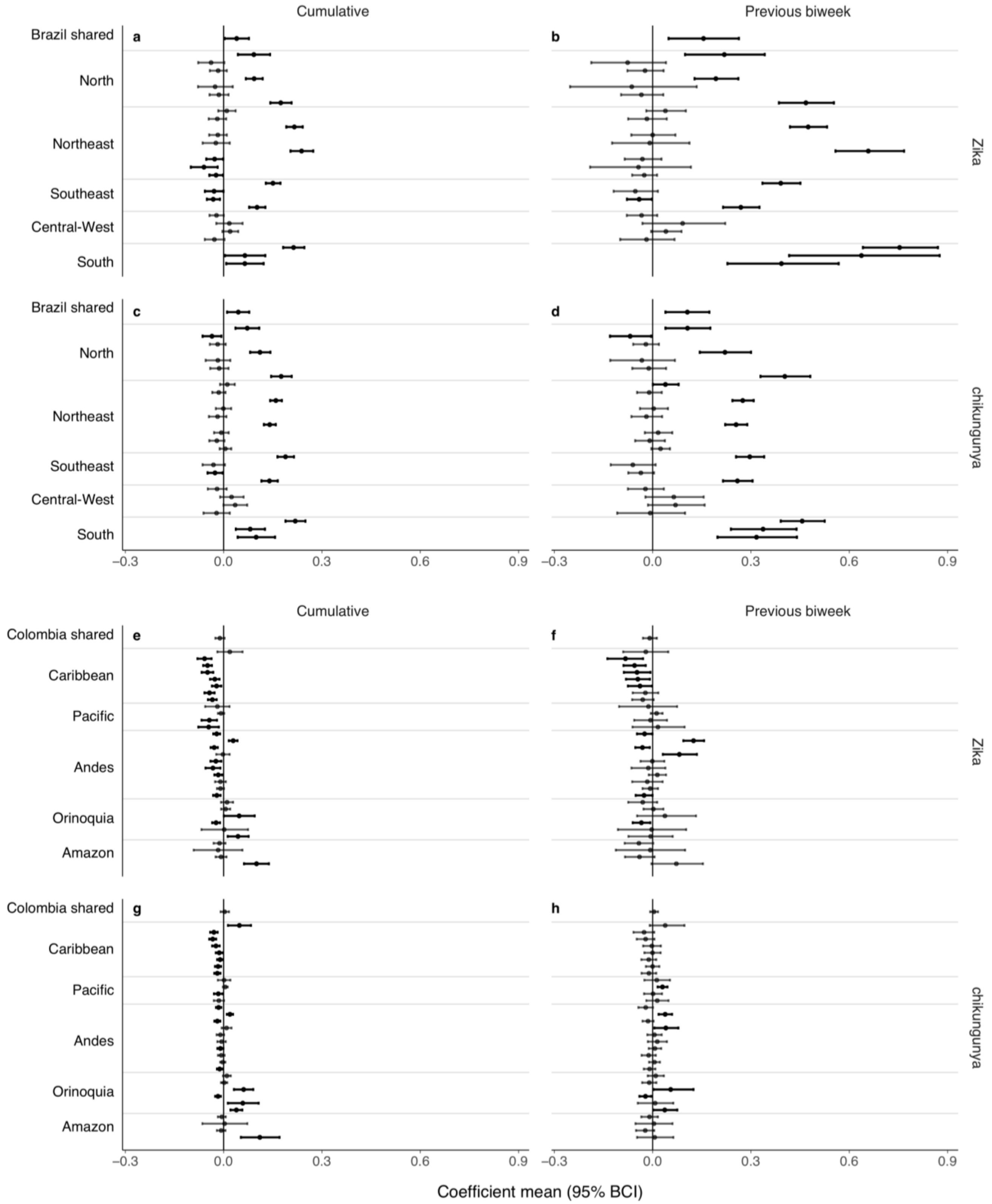

**SI Figure 7:** Correlation between arbovirus related case count totals for states in Brazil (**a, b, d, e**) and departments in Colombia (**c**). Correlation coefficients are displayed in the bottom right corner of each panel.

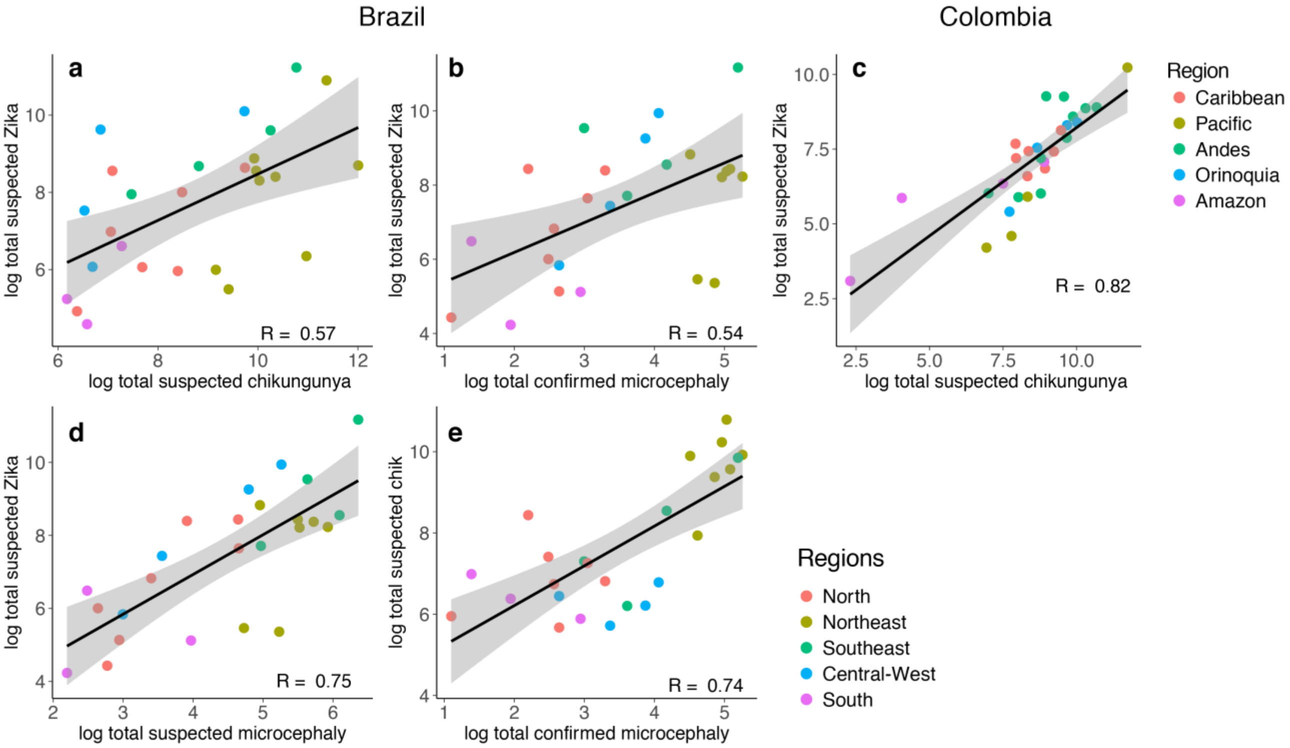

**SI Figure 8:** Hierarchical model shared effects for permuted recent year datasets. Mean and 95% Bayesian credible intervals are displayed for shared effect coefficients from models fit using the actual dataset (leftmost panels) and alternative datasets where 2015 to 2017 where replaced with three consecutive years of data preceding 2015 for Brazil and Colombia. Nonsignificant results are displayed with light shading.

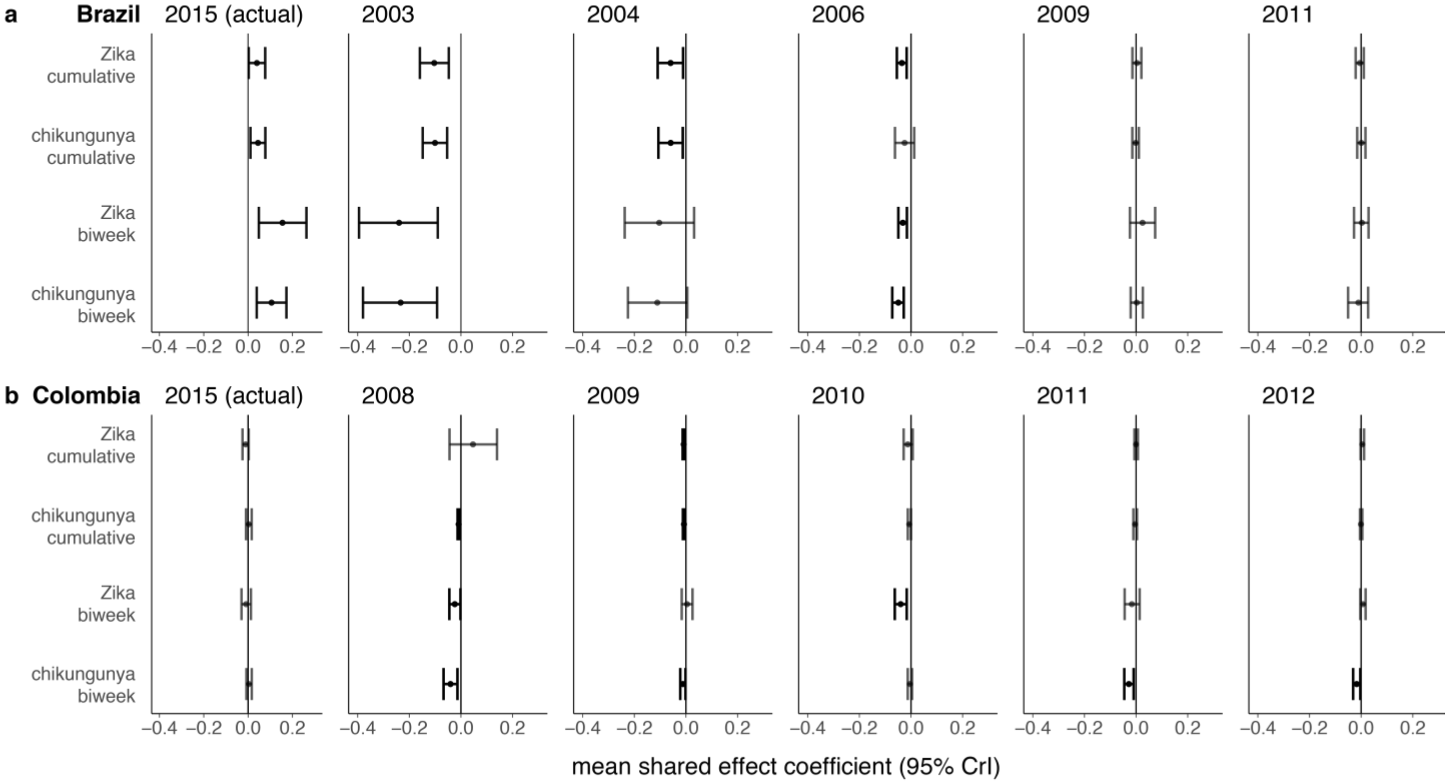

**SI Figure 9:** Effects of immune-mediated interactions between DENV and ZIKV on case counts in stochastic simulations. For each combination of cross-protection, enhancement, and $R_0$ pair, the average ratio (over 100 simulations) between cumulative DENV cases over 1 year (**a**) and 2 years (**b**) after the introduction of ZIKV is plotted against the ratio of cumulative ZIKV cases with and without immune-mediated interactions with DENV. Values above 1 indicate increases in counts and values below 1 indicated decreases compared to the average value from the corresponding scenario without enhancement or ZIKV cross-protection against DENV. For each enhancement and cross-protection scenario pair, linear model fits to the averages (over the 5 $R_0$ pairs considered) are displayed as gray lines. Negative slopes are consistent with the hypothesis that higher ZIKV incidence is associated with lower DENV incidence. For the case when DENV $R_0 = 2$ and ZIKV $R_0 = 2$, panel **c** shows the relative changes in cumulative DENV incidence over 2 years post-ZIKV introduction (blue points) and changes in DENV peak size (red points). Black vertical lines separate the enhancement scenarios and the level of cross-protection increases from left to right within these sections. Relative changes in peak size are based on the highest DENV incidence in 20 years post-ZIKV introduction versus 20 years pre-ZIKV.

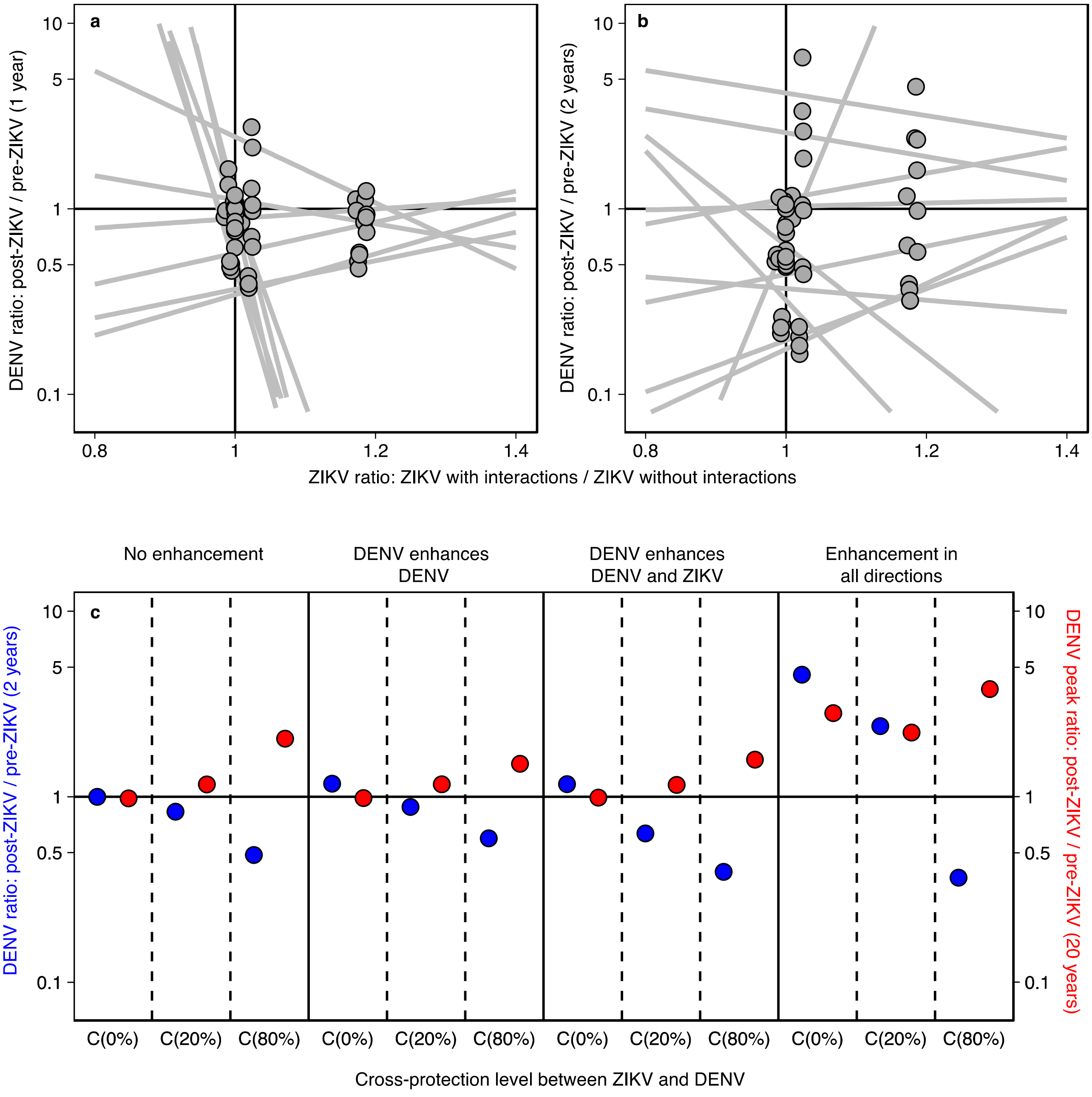
